# Haplotype-resolved genome assembly of upland switchgrass provides insights into cold tolerance

**DOI:** 10.1101/2024.08.26.609807

**Authors:** Bingchao Wu, Dan Luo, Yuesen Yue, Haidong Yan, Min He, Xixi Ma, Bingyu Zhao, Bin Xu, Jie Zhu, Jing Wang, Jiyuan Jia, Min Sun, Zheni Xie, Xiaoshan Wang, Linkai Huang

**Author notes:** these authors contributed equally to this work (Bingchao Wu, Dan Luo). Corresponding author: Linkai Huang.

## Abstract

Switchgrass (*Panicum virgatum* L.) is a bioenergy and forage crop. Upland switchgrass exhibits superior cold tolerance than lowland ecotype, but the underlying molecular mechanisms remain unclear. Here, we presented a high-quality haplotype-resolved genome of the upland ecotype ‘Jingji31’ and conducted multi-omics analysis to understand its cold tolerance. The divergence between upland and lowland ecotypes of switchgrass occurred after the differentiation of the two subgenomes (K and N). Under cold stress, the K subgenome has more differentially expressed genes (DEGs). Transcriptome analysis revealed ecotype-specific differential expressions among members of the cold-responsive (*COR*) gene families. Specifically, certain members of the *AFB1*, *ATL80*, *HOS10*, and *STRS2* gene families exhibited opposite expression changes between the two ecotypes, potentially contributing to their differential cold tolerance. By using haplotype-resolved genome, we identified more cold-induced allele-specific expressions (ASEs) in the upland ecotype, and these ASEs were significantly enriched in the *COR* gene families. Genome-wide association study detected an association signal on Chr3K related to overwintering rate, which overlapped with a selective sweep region and contained a cytochrome P450 (*CYP450*) gene highly expressed under cold stress. Heterologous overexpression of *CYP450* in rice alleviated leaf wilting and improved cold tolerance. Our study provides a high-quality haplotype-resolved genome of upland switchgrass, to advance conceptual understanding of plant cold tolerance for breeding crops with enhanced cold adaptation.

## Introduction

Switchgrass (*P. virgatum*) is a perennial C4 grass utilized as both a forage crop and a dedicated feedstock for bioenergy production^1,2^. It comprises two distinct ecotypes, lowland and upland, displaying substantial variations in morphology and environmental adaptability^3,4^. The lowland ecotype typically thrives in warm, moist environments, featuring greater plant height and broader leaves. In contrast, the upland ecotype predominantly inhabits cold, arid areas and is able to overwinter in colder temperate zones^5^. The upland ecotype likely contains gene resources conferring cold tolerance that differ from those in the lowland ecotype.

Due to the frequent occurrence of extreme weather events caused by global climate change, stable crop production faces significant challenges. Cold stress is one of the most threatening abiotic stresses in the growth and development of plants, impacting the geographical distribution of plants and even leading to plant mortality, thereby causing a decrease in crop yield^6^. Additionally, planting switchgrass on marginal lands is an effective way to increase its cultivation area, but these lands often face various abiotic stresses, especially the threat of low temperature. Upland switchgrass represents an ideal model for understanding how plants respond to cold stress, as it can successfully overwinter in northern cold regions compared to lowland ecotypes. However, few studies have explored the molecular mechanisms underlying the regulation of cold stress responses in the upland ecotype relative to the lowland ecotype, and the underlying mechanisms remain unclear.

For most plants with an open-pollination mechanism, the common strategy for selecting superior germplasm is to transfer the excellent traits of parents to the offspring through hybridization. The improvement of offspring traits is typically caused by increased genetic variation and specific expression of allelic genes at certain loci^7^. The time and space-specific expression of different allelic genes can result in significant differences in gene products and lead to distinct phenotypes^8,9^. For highly heterozygous switchgrass, the previously reported collapsed representation may have overlooked half of the heterozygous variants in the genome and could have introduced assembly errors in regions of divergence between haplotypes^10^.

In this study, we uncover the cold tolerance mechanism of upland switchgrass through constructing a high quality and haplotype-resolved reference genome of upland switchgrass ‘Jingji31’ together with integrating transcriptomics, population genetics, and functional validation assays. We identified a large number of *COR* genes with opposite expression trends in upland and lowland switchgrass under cold stress, which may contribute to the cold tolerance differences between the two ecotypes. Additionally, ASEs were widely present in switchgrass and were induced by cold stress, particularly ASEs of certain *COR* genes. GWAS and selective sweep analysis identified a large number of candidate genes potentially associated with cold tolerance, among which overexpression of one such candidate gene positively regulated cold tolerance. Our findings not only improve the understanding of cold tolerance in upland switchgrass to accelerate genome-assisted breeding of cold tolerance in this important model energy plant, but also hold promise in promoting comparative genomics studies of other crops.

## Results and Discussion

### Assembly and annotation of the upland switchgrass genome

The genome size of the switchgrass upland ecotype ‘Jingji31’ (abbreviated as ‘JJ31’)was estimated to be ∼1.19 Gb using k-mer analysis based on 67.8 Gb (57× coverage) of Illumina short-read data (Fig. 1a, Supplementary Table 1, 2). A total of 53.2 Gb (44.7× coverage) of PacBio high-fidelity long read (HiFi) sequences and 145.2 Gb (122× coverage) Illumina-sequenced Hi-C data were then generated (Supplementary Table 1). Due to the high heterozygosity (1.55%), Hfiasm was used for haplotype-resolved de novo assembly using phased assembly graphs^11^. Two phased haplotypes, ‘JJ31-A’ and ‘JJ31-B’, were anchored to pseudo-chromosomes using Hi-C reads. The Hi-C interaction map demonstrated that our chromosome-level anchoring was of high quality and reliability (Fig. 1b). Furthermore, through co-linearity analysis with the previously published lowland ecotype switchgrass ‘AP13’ genome, all pseudo-chromosomes of the two haplotypes were successfully assigned to different subgenomes, bearing the same chromosome IDs "K" and "N"^10^ (Fig. 1c and Supplementary Fig. 1). Ultimately, the genome sizes of ‘JJ31-A’ and ‘JJ31-B’ were determined to be 1.14 Gb and 1.11 Gb, respectively, with 98.47% (‘JJ31-A’) and 99.02% (‘JJ31-B’) of sequences anchored across 18 chromosomes (Fig. 1d, Table 1, Supplementary Table 3 and 4). Their respective Contig N50 values were 4.9 times and 4.7 times higher than that of the ‘AP13’ genome (5.5 Mb)^10^ (Table 1).

**Fig. 1.**
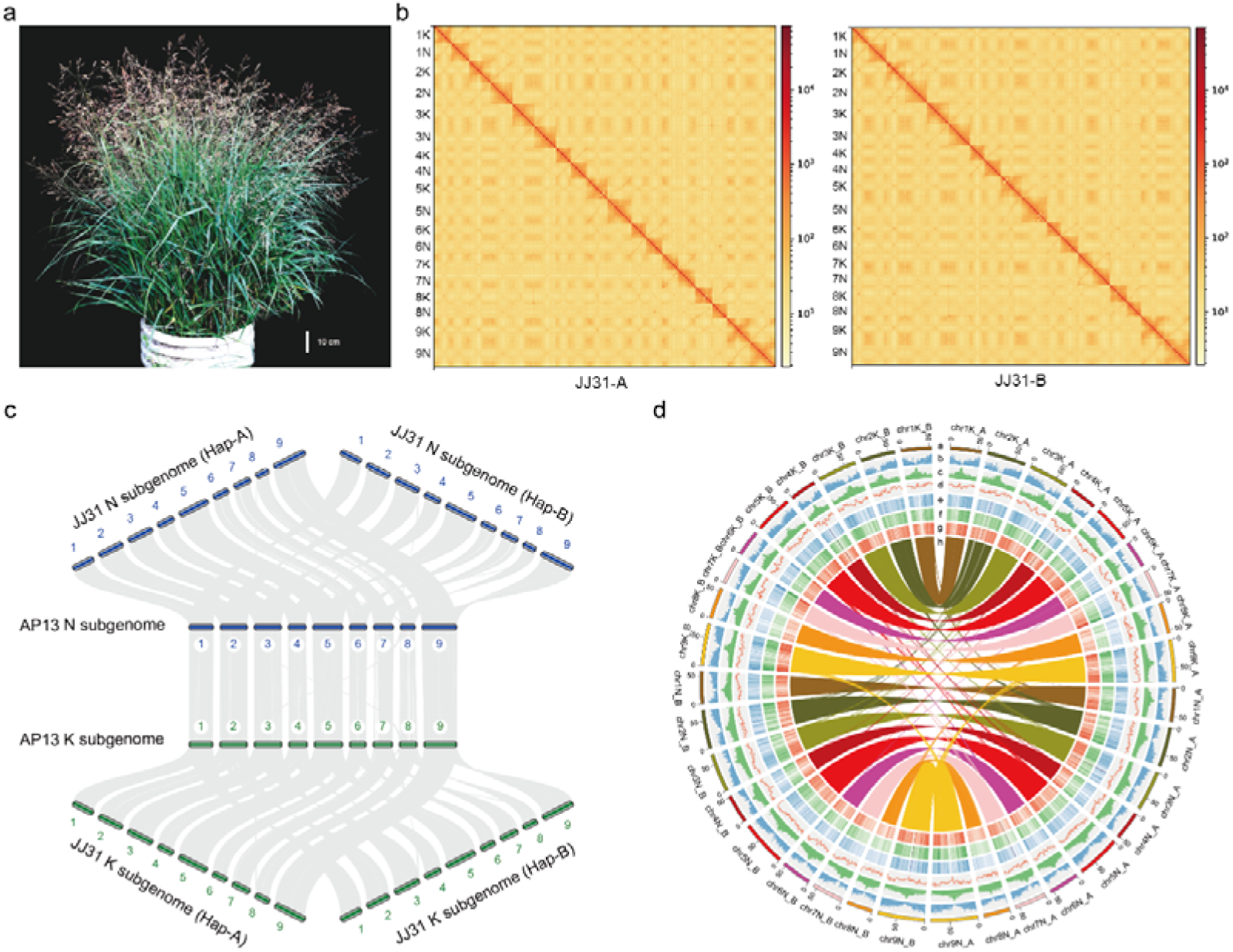
High-quality haplotype-resolved genome assembly of upland switchgrass JJ31. **a**, Flowering morphology of the upland ecotype JJ31. **b**, Whole genome Hi-C heat map for ‘JJ31-A’ (left) and ‘JJ31-B’ (right). **c**, The chromosome collinearity between the genomes of JJ31 and Alamo. Numbers represent chromosome identifiers. **d**, Genome features of the JJ31 genome. Track ‘a’: chromosome length; track ‘b’: gene density; track ‘c’: transposable element (TE) density; track ‘d’: GC content; track ‘e-g’: gene expression levels in roots, stems, and leaves; track ‘h’: collinearity between chromosomes of ‘JJ31-A’ and ‘JJ31-B’.

**Table 1.** Summary of assembly and annotation of two haplotype genomes in upland switchgrass.

|  | JJ31-A | JJ31-B |
| --- | --- | --- |
| Contig N50 (Mb) | 27.05 | 25.98 |
| Contig length (Mb) | 1,124.57 | 1,136.95 |
| Scaffold N50 (Mb) | 67.96 | 62.02 |
| Scaffold length (Mb) | 1,143.31 | 1,110.80 |
| Chromosome anchoring rate (%) | 98.47 | 99.02 |
| Gene no. | 79,672 | 79,416 |
| Repeat sequence length (Mb) | 619.58 | 594.02 |
| Repeat ratio (%) | 54.19 | 53.48 |
| QV | 45.16 | 50.27 |

We employed various strategies to assess the quality of the haplotype-resolved genomes. We realigned paired-end reads to the two haploid genomes, resulting in observed alignment rates of 98.07% and 97.93%, respectively (Supplementary Table 3). Furthermore, the embryophyta Benchmarking Universal Single-Copy Orthologs (BUSCO) analysis indicated completeness rates of 98.4% and 98.1% for the two haplotypes (Supplementary Table 3). Using Merqury to calculate quality values (QV), ‘JJ31-A’ and ‘JJ31-B’ achieved values of 45.16 and 50.27, respectively, exceeding the Vertebrate Genomes Project standard of QV40^12^ (Table 1). The long terminal repeat (LTR) assembly index (LAI) for both phased haplotype genomes approached 20, nearly meeting the gold standard^13^ (Supplementary Table 3). These results affirm the accuracy, completeness, and contiguity of our two haploid genome assemblies.

Both haplotype-resolved genomes were annotated with a comprehensive strategy that combined homolog prediction, de novo prediction, and other evidence-driven predictions. In the two haplotypes, 79,672 and 79,416 protein-coding genes were predicted, with 98.9% supported by known gene function databases (Table 1 and Supplementary Table 3). The gene models of the two haploid genomes have an average coding sequence length of ∼1 kb and an average of four exons per gene (Supplementary Table 3). We also identified 0.62 and 0.59 Gb repeat sequences, accounting for 54.19% and 53.48% of the two haplotypes, respectively, of which 42.09% and 41.51% are LTRs (Supplementary Table 3). Additionally, we identified 2,245 and 2,206 miRNAs, 1,264 and 1,291 tRNAs, 9,171 and 5,221 rRNAs, as well as 1,190 and 1,154 snRNAs in the two haplotype genomes, respectively (Supplementary Table 3). Considering the overall superior quality of the ‘JJ31-B’ haplotype, it was selected for subsequent analysis unless otherwise stated.

### Enhanced cold tolerance and subgenome dominance in upland switchgrass indicated by comparative genomics analysis

To understand the evolutionary relationship between the two ecotypes of switchgrass, we performed comparative genomic analysis by adding several closely related species. Based on the phylogenetic tree, we found that the K and N subgenomes of switchgrass diverged approximately 6.6 million years ago (Mya), while the divergence between switchgrass and *P. hallii* occurred around 8.8 Mya, consistent with previous studies^10^ (Fig. 2a). It is noteworthy that the K subgenomes of the two ecotypes diverged at 2.3 Mya, and the N subgenomes diverged at 2.6 Mya (Fig. 2a), suggesting that the divergence between these two ecotypes may have occurred during this period. We further determined the divergence time between the two ecotypes to be ∼2.2 Mya (Ks peak at 0.03) by calculating the synonymous substitution rate (Ks) of orthologous gene sets between the different subgenomes of the two ecotypes (Fig. 2b and Supplementary Table 5). Combining previous report on the tetraploidization time of switchgrass (< 4.6 Mya)^10^, we hypothesize that the evolutionary timeline of switchgrass first involved the differentiation of the N and K subgenomes, followed by tetraploidization, and then the differentiation of the two ecotypes.

**Fig. 2.**
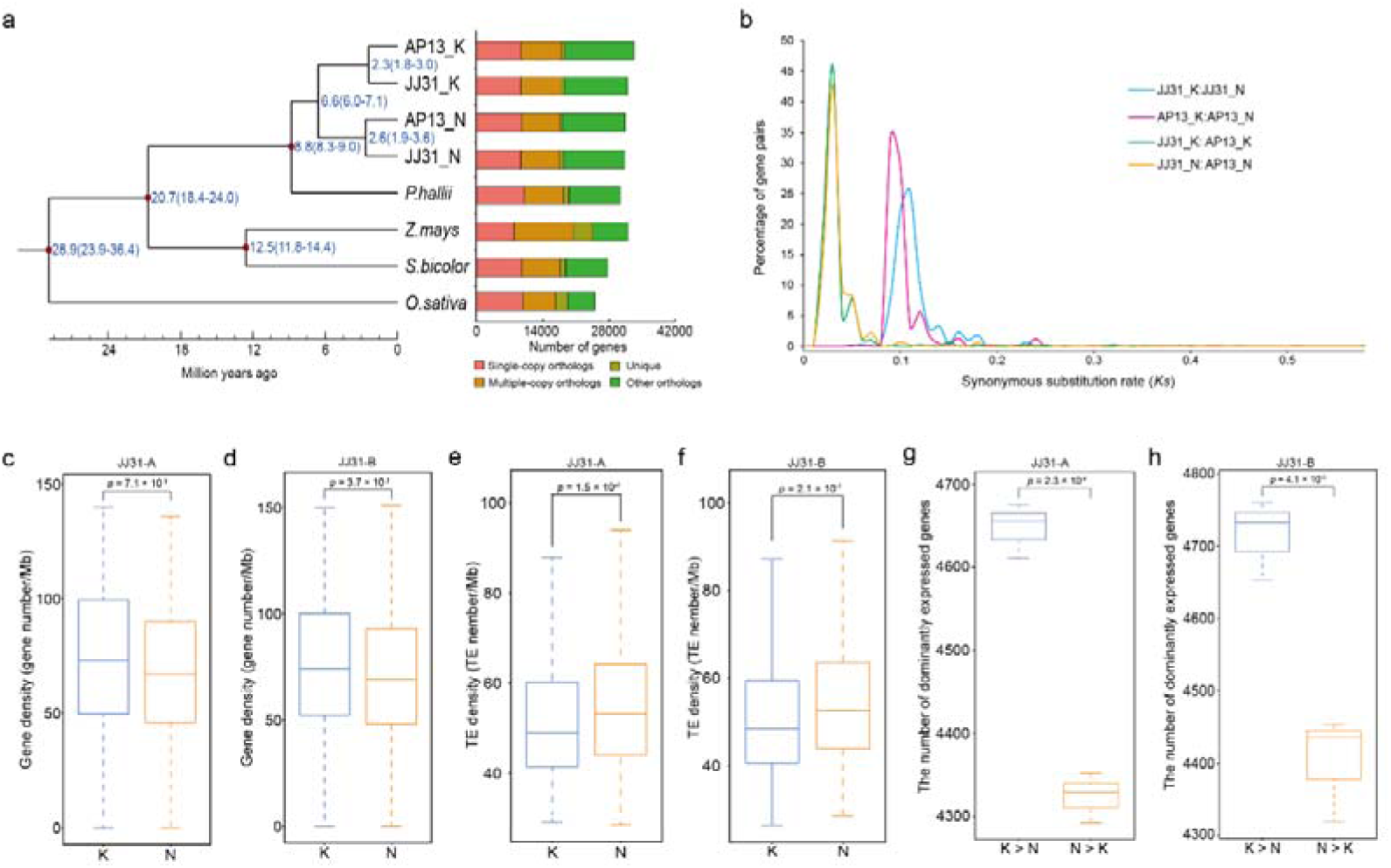
Comparative genomics and subgenome dominance analysis. **a**, Phylogenetic tree and divergence time estimates of ‘JJ31’ and five closely related species. The right panel shows the distribution of single-copy, multi-copy, unique, and other gene orthologs. **b**, Evolutionary analysis of the ‘JJ31’ and ‘AP13’. The Ks distribution is shown for orthologs in the switchgrass genomes. K and N represent two subgenomes, respectively. **c**, **d**, Gene density of the two subgenomes in the ‘JJ31-A’ (c) and ‘JJ31-B’ (d). **e**, **f**, TE density of the two subgenomes in the ‘JJ31-A’ (c) and ‘JJ31-B’ (d). **g**, **h**, Number of dominantly expressed genes in the two subgenomes of the ‘JJ31-A’ (g) and ‘JJ31-B’ (h). The statistical window size is 1 Mb.

It was reported that upland switchgrass exhibits better cold tolerance than the lowland ecotype^14^. Compared to the ‘AP13’, 4,147 expanded, 5,030 positively selected, and 4,873 specific genes were identified in ‘JJ31’. These genes were enriched in several stress-regulated pathways and biological processes (Supplementary Fig. 2). The expanded and specific genes in ‘JJ31’ were mainly enriched in calcium channel activity, calcium ion transmembrane transporter activity, melatonin receptor activity, and sphingolipid metabolic pathways, which are believed to play crucial roles in protecting plants from cold stress^15–19^. These positively selected specific genes were associated with G-protein coupled receptor activity, which was involved in signal transduction of plant stress responses^20^. The above results may be one of the reasons leading to the greater cold tolerance of upland switchgrass compared to the lowland ecotype.

Subgenome dominance is a common phenomenon in polyploid plants, prompting our investigation into the subgenome characteristics within the two haplotypes of upland switchgrass. We employed a sliding window approach (window size 1 Mb) to assess gene density and transposable element (TE) density in the two subgenomes. Compared to the N subgenomes in ‘JJ31-A’ and ‘JJ31-B’, the K subgenomes exhibited higher gene density (71.2 versus 66.2 genes per Mb and 73 versus 67.6 genes per Mb, *P* < 0.01), more genes with dominant expression (4,662 versus 4,305 and 4,715 versus 4,403, *P* < 0.01), less TE density (53.5 versus 56.4 TEs per Mb and 52.5 versus 55.7 TEs per Mb, *P* < 0.01) (Fig. 2c-h and Supplementary Tables 6-9). Additionally, the cold stress transcriptome data from both ecotypes revealed slightly more DEGs on the K subgenome, suggesting that it might play a more significant role in response to cold stress (Supplementary Tables 10). In summary, all our statistics regarding the two subgenomes pointed to the K subgenome as the dominant one in both haplotypes, consistent with the findings in the ‘AP13’ genome^10^.

### Contribution of specific differential expression of *COR* gene families to the cold tolerance differences between the two ecotypes

The upland ecotype ‘JJ31’ exhibited greater cold tolerance than the lowland ecotype ‘Alamo’ (noting that ‘AP13’ was a line/clone selected from ‘Alamo’)^14^ based on the phenotypic and physiological indicators under cold stress (Fig. 3a-c). Leaves of both ecotypes showed significant wilting after 56 days of cold treatment, but only ‘JJ31’ regrew new leaves after returning to room temperature (Fig. 3a). In addition, relative water content (RWC), relative electrical conductivity (REC), and malondialdehyde (MDA) showed significant changes (*P* < 0.05) in ‘JJ31’ after 21 days of cold stress compared to the control, while in ‘Alamo’, RWC significantly decreased and REC and MDA significantly increased after 14 days of cold stress compared to the control (*P* < 0.05) (Fig. 3b,c). The slower physiological changes under cold stress might indicate better cold tolerance in ‘JJ31’ than in ‘Alamo’.

**Fig. 3.**
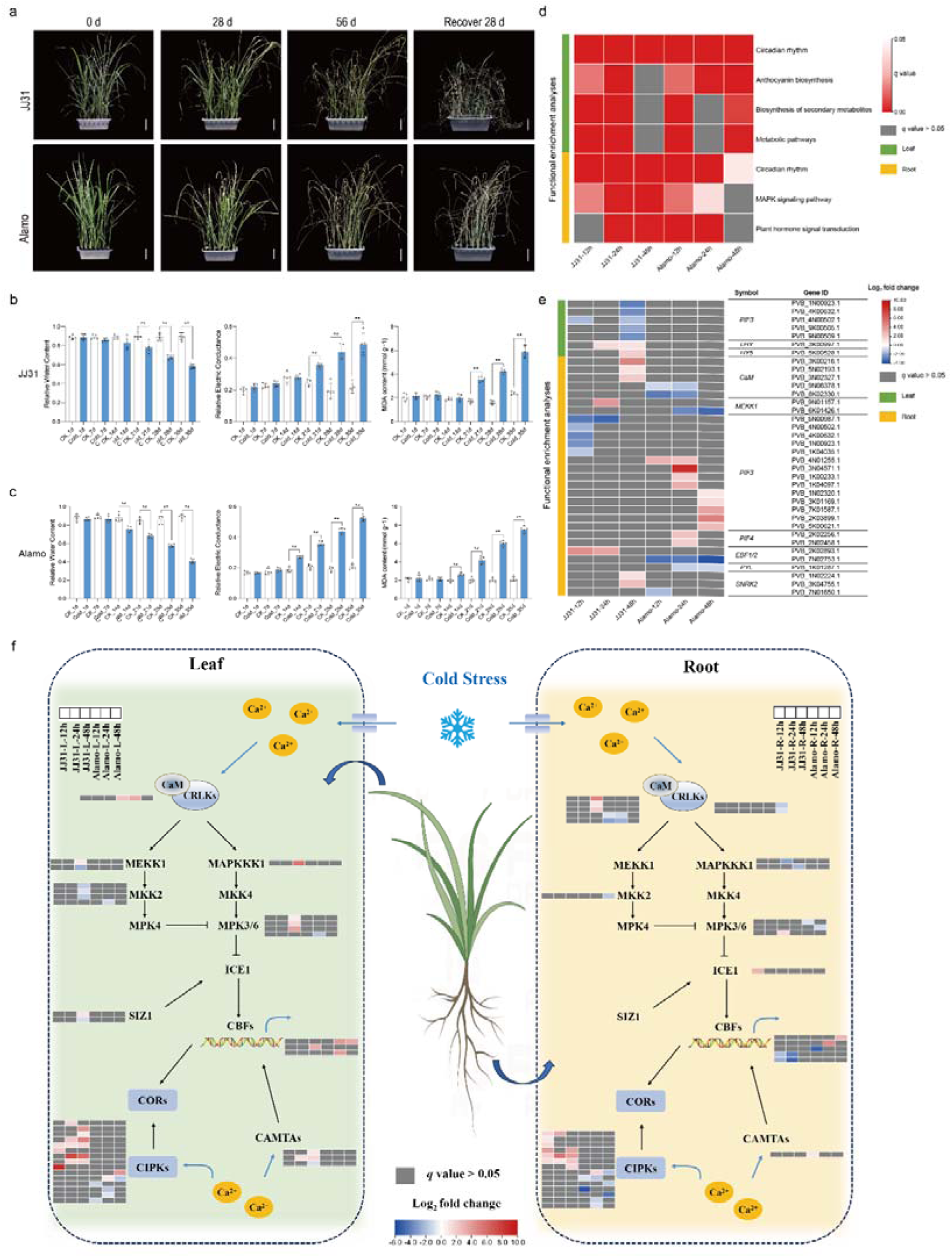
Transcriptional landscape differences between the two ecotypes under cold stress. **a**, Phenotypic changes of JJ31 and Alamo on days 0, 28, and 56 under cold stress at 4 °C and recover at room temperature for 28 days. Scale bar indicates 7 cm. **b**, Changes in physiological indicators of JJ31 under control (room temperature) and cold stress (4 °C). From left to right: RWC, REC, and MDA. ** indicates *P* < 0.005. **c**, Changes in physiological indicators of Alamo under control (room temperature) and cold stress (4 °C). From left to right: RWC, REC, and MDA. ** indicates *P* < 0.005. **d**, KEGG enrichment of all compared DEGs identified in leaves and roots of JJ31 and Alamo under cold stress. **e**, Expression changes of cold-tolerance-related genes in the circadian rhythm, MAPK signaling pathway, and plant hormone signaling transduction pathways in two ecotypes. **f**, Expression changes of key genes in the *CBF*-dependent cold response pathway in the two ecotypes.

To understand the molecular response mechanisms underlying the greater cold tolerance of upland switchgrass, we conducted transcriptome sequencing on the leaves and roots of ‘JJ31’ and ‘Alamo’ at three time points under cold stress (Supplementary Table 1). Based on KEGG enrichment analysis, the DEGs revealed in each comparison group were mainly enriched in circadian rhythm, MAPK signaling pathway, and plant hormone signal transduction pathways (Fig. 3d and Supplementary Fig. 3). Although both ecotypes relied on similar pathways to respond to cold stress, we found that genes related to cold tolerance within these pathways exhibited ecotype-specific expression (Fig. 3e). During the initial exposure of plants to cold stress, Ca^2+^ influx induces activity of calmodulin (CaM) and activates downstream MEKK1 to positively regulate plant cold tolerance^21,22^. Our study found that three members of the *CaM* family and one member of the *MEKK1* family were specifically upregulated in ‘JJ31’, while two and one members of the respective families were specifically downregulated in ‘Alamo’ (Fig. 3e). Two transcription factors, PIF3 and PIF4, are reported to negatively regulate plant cold tolerance by inhibiting the expression of *CBF*, while the EIN3-BINDING F-BOX 1/2 (EBF1/2) proteins enhance cold tolerance by degrading PIF3^23,24^. We found that nine *PIF3* genes were specifically downregulated in ‘JJ31’, while nine *PIF3* and two *PIF4* genes were specifically upregulated in ‘Alamo’, with two *EBF1/2* genes exhibiting opposite expression changes (Fig. 3e). Additionally, we found that one *LHY*, one *HY5*, and two *SNRK2* genes were specifically upregulated in ‘JJ31’, while one *SNRK2* and one *PYL* were specifically downregulated in ‘Alamo’ (Fig. 3e). These genes have been reported to positively regulate plant cold tolerance^25–28^. These results suggested that the ecotype-specific expression of these cold stress regulatory genes might interpret differences in cold tolerance between the two ecotypes.

The above results imply that there may be more cold-response (*COR*) genes with ecotype-specific expression. We identified 795 *COR* genes in switchgrass based on the 109 *COR* gene families reported in *Arabidopsis*^29^ (Supplementary Table 11). We identified 182 and 251 specific DEGs significantly enriched in *COR* genes in ‘JJ31’ and ‘Alamo’, respectively (*P* = 0.035 and *P* = 1.0745e-5, Supplementary Fig 4, Supplementary Table 12, 13 and Supplementary Note 1). These *COR* genes belonged to 58 and 55 families, respectively, among which members of 25 families showed specifically differential expression in the leaves or roots of only one ecotype (Supplementary Fig. 5a). The members of 44 *COR* gene families exhibited differential expression in both ecotypes, but we noted that some of these genes showed opposite expression changes between the two ecotypes (Supplementary Fig. 5b).

The auxin signaling F-box protein 1 (AFB1) -mediated auxin signaling pathway is involved in plant tolerance to abiotic stresses, and its overexpression can enhance plant tolerance to salt and cold stresses^30^. Three *AFB1* genes were upregulated in ‘JJ31’, while four *AFB1* genes were downregulated in ‘Alamo’ (Supplementary Fig. 5b). ATL80, an E3 ubiquitin ligase and negative regulator in response to cold stress^31^, had three genes downregulated in ‘JJ31’ and one gene upregulated in ‘Alamo’ (Supplementary Fig. 5b). The *Arabidopsis* mutant *hos10-1* was reported to be completely unable to acclimate to the cold^32^. We found that 15 *HOS10* genes were upregulated in ‘JJ31’, while 11 *HOS10* genes were downregulated in ‘Alamo’ (Supplementary Fig. 5b). Interestingly, although STRESS RESPONSE SUPPRESSOR2 (STRS2) was reported to negatively regulate *Arabidopsis* tolerance to salt, osmotic, and heat stress, and not cold stress^33^, our study found that three *STRS2* genes were downregulated in ‘JJ31’ and two were upregulated in ‘Alamo’ (Supplementary Fig. 5b), which may highlight the role of STRS2 in response to cold stress in switchgrass. In summary, the ecotype-specific expression of the above *COR* genes or their opposite expression changes in the two ecotypes might contribute to their differences in cold tolerance.

The transcriptional regulatory pathway dependent on *CBF* is crucial for plant response to cold stress^34–36^. Similarly, We found that cold response genes in *CBF*-dependent pathway was activated to varying degrees in both ‘JJ31’ and ‘Alamo’, with slight differences between leaves and roots (Fig. 3f). We observed that the specific differential expression of *MEKK1* and *SIZ1* genes occurred only in the leaves, while the specific differential expression of *CRLKs* and *ICE1* genes occurred only in the roots (Fig. 3f). Additionally, members of the *CaM*, *MPK3/6*, and *CIPKs* families tended to exhibit opposite expression trends between the two ecotypes (Fig. 3f). In conclusion, through the identification of *COR* gene families and comparative transcriptome analysis, we comprehensively revealed the landscape of differential expression of *COR* genes between the two ecotypes, which may contribute to the superior cold tolerance of the upland compared to the lowland ecotype.

### The involvement of ASE in response to cold stress revealed by haplotype-resolved genome

The haplotype-resolved genome of upland switchgrass enabled us to use RNA-seq data to identify ASEs, which have been reported in recent studies to profoundly impact plant growth and development^37^. According to the correlation between the number of ASEs and the number of transcriptome samples used in the analysis, we aimed to obtain a complete ASE collection in switchgrass using sufficient data. We found that the number of ASEs stabilized when the number of transcriptome samples reached 15 by utilizing transcriptome data reported by Zuo et al (with at least 2 replicates)^38^(Fig. 4a). A total of 16,801 ASEs were identified in switchgrass, with significantly more ASEs biased towards ‘JJ31-A’ expression than towards ‘JJ31-B’ (Fig. 4b and Supplementary Table 14). To understand the impact of natural selection on ASEs and non-ASEs, we calculated the Ka/Ks ratio between allele pairs. Although most allele pairs exhibited low Ka and Ks values, ASEs underwent significantly stronger purifying selection pressure compared to non-ASEs (Fig. 4c and Supplementary Table 15). To further investigate potential causes of ASE, we examined the distribution patterns of SNPs surrounding ASEs and equivalently expressed alleles (EEAs). Compared to EEAs, ASEs exhibited significantly higher SNP density in the upstream, exonic, intronic, and downstream regions, as with previous findings in other plants^39^ (Fig. 4d). The SNP density in the upstream region was higher than that in other regions, suggesting that the occurrence of ASE might correlate with the greater variation in the upstream region of the gene.

**Fig. 4.**
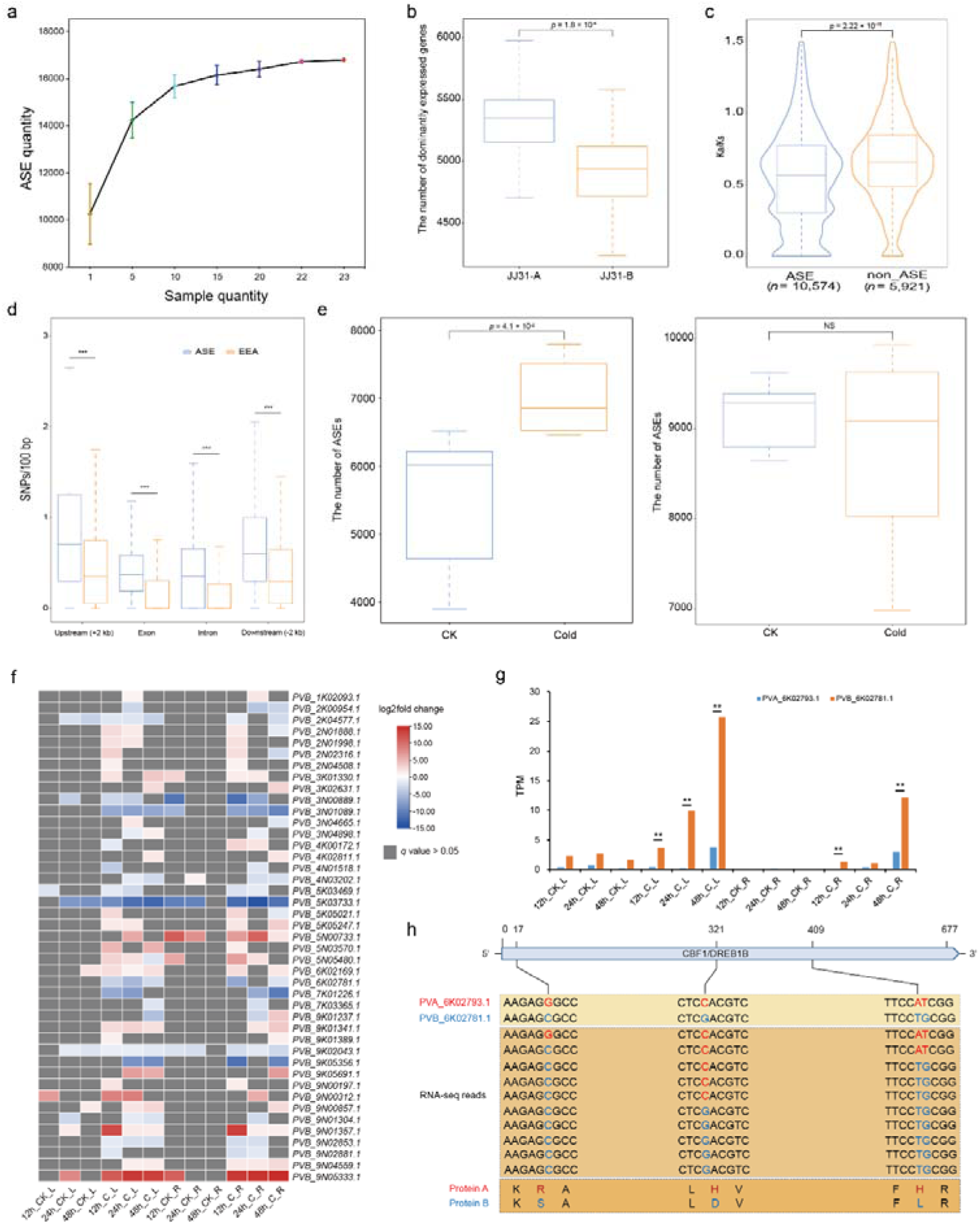
ASEs in switchgrass. **a**, ASE numbers increase with the quantity of RNA-seq samples. The specified number sets were selected randomly from 23 ASE sets with three replicates. **b**, Dominant expressed alleles in two haplotype genomes. **c**, *Ka/Ks* of ASE and non-ASE genes. Minima and maxima are present in the lower and upper bounds of the whiskers, respectively, and the width of violin are densities of *Ka/Ks* value. *P* values were calculated with two-sided Student’s t-test. **d**, SNP density in ASE and EEA genes. The y axis represents SNP numbers every 100 bp. *P* values were calculated with two-sided Student’s t-test. *** indicates *P* < 0.0005. **e**, Number of ASE genes identified in JJ31 (left) and Alamo (right) under control (room temperature) and cold stress (4 °C). *P* values were calculated with two-sided Student’s t-test. NS indicates not significant. **f**, The expression changes of alleles of 43 *COR* genes in JJ31 under control (room temperature) and cold stress (4 °C) conditions. ASE of these genes was induced by cold stress at least at one time point. **g**, The expression levels (TPM) of two alleles of the *CBF* gene (*PVA_6K02793.1* and *PVB_6K02781.1*) across different transcriptome samples. ** indicates adjusted *P*-value < 0.01. **h**, Pattern diagram of *PVB_6K02781.1* advantage expression. Red indicates the allele ID and the corresponding bases and encoded amino acid types in ‘JJ31-A’; blue indicates the allele ID and the corresponding bases and encoded amino acid types in ‘JJ31-B’. "RNA-seq reads" represents the proportion of reads containing different SNP types that map to ‘JJ31-B’. From left to right, the RNA-seq reads aligned to the first SNP site are 79, with 92% supporting C and 8% supporting G; the RNA-seq reads aligned to the second SNP site are 100, with 45% supporting G and 55% supporting C; the RNA-seq reads aligned to the third and fourth SNP sites are both 67, with 81% and 85% supporting T and G, respectively, and 19% and 13% supporting A and T, respectively.

Similarly, we utilized the transcriptome data obtained in this study under cold stress to identify ASEs, aiming to explore whether the response of ASE to cold stress differs between the two ecotypes (Supplementary Note 2). Compared to the control group, a significant increase in ASEs were detected in ‘JJ31’ after experiencing cold stress, while no significant change occurred in ‘Alamo’, indicating that more ASEs in ‘JJ31’ were induced by cold stress (Fig. 4e, Supplementary Table 16, 17). Finally, we identified ‘2,620’ and ‘751’ cold-induced ASEs in two tissues of ‘JJ31’ and ‘Alamo’, respectively (Supplementary Fig 6, 7, Supplementary Table 18 and Supplementary Note 2).

We further found that 43 cold-induced ASEs were significantly enriched in *COR* genes in ‘JJ31’ (*P* = 0.0013, Fig. 4f), while there was no significant enrichment in ‘Alamo’ (*P* = 0.0783), supporting the importance of ASEs in responding to cold stress in ‘JJ31’. Among these genes, we observed that two alleles of a well-known *CBF* gene, *PVA_6K02793.1* and *PVB_6K02781.1*, did not exhibit differential expression in the control group, but *PVB_6K02781.1* showed significant preferential expression in response to cold stress (Fig. 4g). Although the sequence similarity between the two alleles is as high as 98.82%, four SNPs cause changes in three amino acids (Supplementary Fig. 8). We aligned the RNA-seq data to the reference genome ‘JJ31-B’ and found a higher proportion of reads containing SNPs corresponding to the B allele type, supporting the dominant expression of *PVB_6K02781.1* (Fig. 4h). In conclusion, our findings indicated the widespread presence of ASE phenomena in switchgrass, and more ASEs were involved in the response to cold stress in upland switchgrass.

### Identification of genes associated with cold tolerance by population genetic analysis

To explore cold tolerance genes in upland switchgrass at the population level, we aligned resequencing data from 340 accessions (242 upland and 98 lowland) reported previously to the ‘JJ31’ genome^10^, resulting in 10,654,902 SNPs and 243,831 SVs (Supplementary Fig. 9, Supplementary Table 19 and Supplementary Note 3). A total of 103.7 Mb and 125.9 Mb of genomic sequences covering 5,084 and 8,428 genes were detected using the sliding window method based on SNPs and SVs, respectively (Supplementary Fig. 10). We found that 66 and 111 genes from the two datasets were significantly (*P* = 0.0196 and *P* = 0.0018, respectively) and annotated as belonging to the *COR* gene family (Supplementary Fig. 10). Among the *COR* genes identified based on SNPs and SVs, approximately 54.5% (36 out of 66) and 62.2% (69 out of 111), respectively, showed differential expression under cold stress (Supplementary Table 20). These results suggested that the differential selection of certain *COR* genes in the two ecotypes potentially contribute to the differences in cold tolerance.

To identify candidate genes related to cold tolerance in switchgrass, we performed a genome-wide association study (GWAS) on the overwintering rate of 340 switchgrass accessions reported previously using two variant datasets^10^ (Supplementary Fig. 11, 12 and Supplementary Table 21, 22). An association signal on Chr3K was simultaneously detected in both SV-GWAS and SNP-GWAS, including an overlapping region with the selective sweep region (Fig. 5a). We examined the expression of 14 genes around the association signal and found that only *PVB_3K03605.1* and *PVB_3K03611.1* were highly expressed under cold stress (Fig. 5b and Supplementary Fig. 13). Interestingly, only *PVB_3K03611.1* appeared in the overlapping region, which encodes cinnamate-4-hydroxylase belonging to the *CYP450* gene family.

**Fig. 5.**
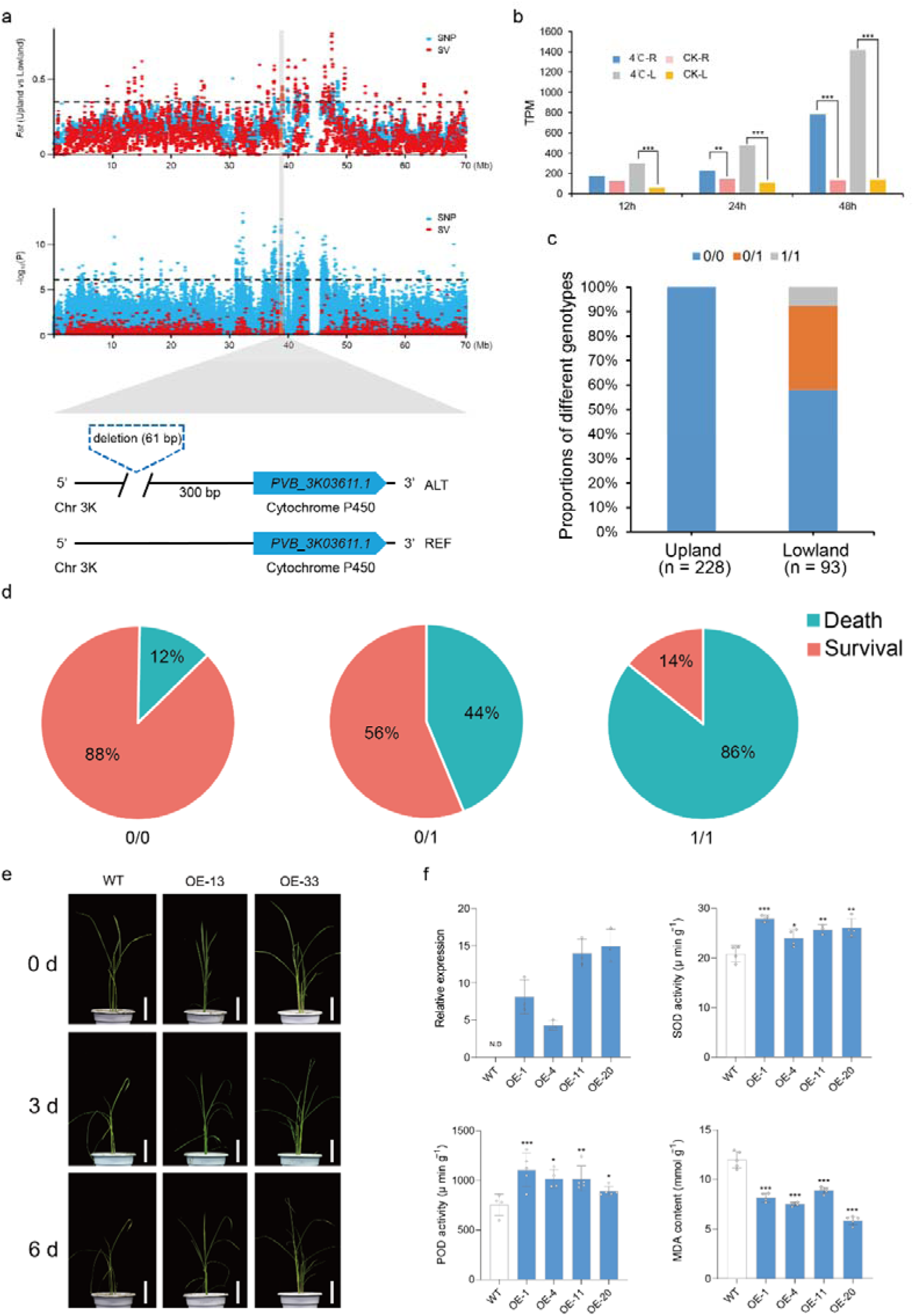
Selective sweep analysis between the two ecotypes and GWAS analysis of overwintering rate. **a**, Upper, selective sweep detection between the two ecotypes based on SNPs and SVs on chromosome 3K using the *Fst* method. The black dashed line represents a cutoff window in which the top 5% data points were selected as sweep regions. Middle, GWAS analysis of overwintering rate based on SNPs and SVs on chromosome 3K. The black dashed line represents the significance threshold based on −log_10_(*P*) > 6. The gray bars represent the overlapping regions between selective sweeps and GWAS. Bottom, schematic diagram of *PVB_3K03611.1* and its upstream 61-bp deletion. **b**, The expression levels (TPM) of *PVB_3K03611.1* under control and cold stress conditions suggest that this gene may potentially positively regulate cold tolerance. *** indicates adjusted *P*-value < 0.001. **c**, The distribution proportions of three genotypes with a 61-bp deletion across different ecotypes of germplasm. 0/0 means consistent with the reference genome, 0/1 means heterozygous, and 1/1 means homozygous. **d**, Overwintering survival rate of three genotype accessions in BRKG area. **e**, Phenotypic changes of rice wild type and overexpression lines on days 0, 3, and 6 under 4°C cold stress, scale bar represents 7 cm. **f**, Physiological parameters of *PVB_3K03611.1* overexpression lines and WT under cold stress. Upper left, relative expression levels of *PVB_3K03611.1* in WT and overexpression lines. N.D indicates not detected; the remaining three figures depict the activities of SOD and POD, as well as the MDA content in WT and overexpressing rice lines after 24 h of cold stress at 4°C. * indicates *P* < 0.05, ** indicates *P* < 0.01, and *** indicates *P* < 0.005.

We found a 61-bp deletion located 300 bp upstream of the promoter region of *PVB_3K03611.1* (Fig. 5a). We further observed this deletion with frequency differences between the two ecotypes, where the 0/1 (heterozygous) and 1/1 (homozygous) genotypes were present in about 40% of the lowland accessions, while the deletion was absent in the upland accessions (Fig. 5c). By analyzing the overwintering rate, it was found that only 12% of germplasms with the 0/0 (same as the reference) genotype failed to overwinter, while the proportions of germplasms with the 0/1 and 1/1 genotypes unable to overwinter were 44% and 86%, respectively (Fig. 5d). These results suggested that this deletion was probably under positive selection in lowland accessions compared to upland accessions and may be potentially associated with cold tolerance.

To validate the role of *PVB_3K03611.1* in cold tolerance, we overexpressed this gene in rice for cold tolerance determination. Compared to the wild type (WT) rice, the transgenic lines exhibited less leaf withering under cold stress (Fig. 5e). Additionally, the transgenic lines displayed significantly higher activities of superoxide dismutase (SOD) and peroxidase (POD), as well as significantly lower levels of MDA than those in WT plants when exposed to low temperatures, indicating that overexpression of *PVB_3K03611.1* enhances the cold tolerance of the transgenic lines (Fig. 5f). Collectively, these results supported that the deletion in the promoter region of *PVB_3K03611.1* might lead to the less cold tolerance trait in the lowland ecotypes than the upland ones.

## Methods

### Sample collection and DNA sequencing

The upland switchgrass cultivar ‘JJ31’ was propagated asexually and planted in three pots in the greenhouse and grown at 26/22 °C (day/night) with photoperiod of 14/10 h of light/dark. Leaves of plants grown at the E3 stage^40^ were collected and pooled for DNA extraction using the DNAsecure Plant Kit (TIANGEN). For Illumina short-reads sequencing, ∼1.5 μg of genomic DNA was extracted to construct a short insert (350 bp) library using a TruSeq Nano DNA HT Sample Preparation Kit. Sequencing was performed using Illumina HiSeq2500 platforms. The raw reads were trimmed using Trimmomatic (v.0.36)^41^ with default parameters. For PacBio HiFi sequencing, SMRTbell libraries were constructed using the SMRTbell Express Template Prep Kit 2.0 (PacBio, CA). Two single-molecule real-time (SMRT) cells were run on the PacBio Sequel II platform. The raw data were processed with the SMRT Link (v.9.0) to obtain HiFi reads, using the parameters --min-passes=3 and --min-rq=0.99. For Hi-C sequencing, the library construction method was the same to the protocol previously used in our laboratory^42^. The constructed library was sequenced using the Illumina NovaSeq 6000 platform.

### Genome size prospection

To estimate the genome size, k-mer analysis (K=17) was performed on Illumina short reads using Jellyfish (v.2.3.0)^43^. The genome size, heterozygosity, and repeat proportion were estimated by GenomeScope (v.2.0)^44^ based on the k-mer frequency distribution. The principle for calculating genome size is based on the formula:*G* = (*N* × (*L* - *k* + 1) - *B*) / *D*, where *N* is the total number of sequence reads, *L* is the average length of the reads, *K* is the k-mer length, *B* is the total number of low-frequency k-mers, *D* is the estimated total depth based on k-mer distribution, and *G* is the genome size.

### Genome assembly and pseudochromosome construction

HiFi reads were assembled into two haplotype-resolved draft genomes using the Hifiasm software (v.0.15.5)^11^. Initially, an all-vs-all pairwise comparison of HiFi reads was performed for self-correction. After haplotype-aware error correction, the corrected reads were used to construct an assembly graph and generate bubbles within this graph. An initial contig assembly based on the overlap graph was obtained using a modified “best overlap graph” strategy. During the assembly process, optimized parameters suitable for polyploid genomes (--n-hap 4) were added to preserve haplotype information as much as possible. Filtered Hi-C reads were aligned to the initial contig assembly using BWA (v.0.7.8)^45^, and the alignment results were used as the input in Juicer (v.1.6)^46^. The 3D-DNA workflow selected only uniquely aligned and valid paired-end reads for further assembly^47^. Finally, the order of scaffolds was manually adjusted using Juicebox (v.2.13.07)^48^ to obtain the final chromosome assembly. HiCExplore (v.3.7.2)^49^ was used to draw heatmaps of the connections between chromosomes.

### Genome assessment

To assess the quality of the genome assembly for accuracy, completeness, and continuity, we used BWA (v.0.7.8)^45^ to map high-quality Illumina paired-end reads to the genome, evaluating the alignment rate and coverage. BUSCO (v.4.1.2)^50^ and the CEGMA (v.2.5)^51^ were used to check the completeness of the genome assembly or annotation. The quality of the genome was further assessed by calculating the QV values with Merqury (v.1.3)^52^ and the LAI with LTR_retriever (v.2.9.8)^13^.

### Annotation of repetitive sequences

We annotated the repetitive sequences by combining homology-based alignment and de novo prediction. The homology-based alignment method used RepeatMasker (v.4.0.5)^53^ and RepeatProteinMask (v.4.0.5)^53^ to identify sequences similar to known repetitive sequences based on the RepBase database (http://www.girinst.org/repbase)^54^. The de novo prediction method utilized LTR_FINDER (v.1.0.7)^55^, Piler (v.3.3.0)^56^, RepeatScout (v.1.0.5)^57^, and RepeatModeler (v.1.0.8)^58^ to construct a de novo repeat sequence library, followed by the use of RepeatMasker (v.4.0.5)^53^ to predict the repetitive sequences in this library.

### Prediction of gene structure

The gene structure was annotated by integrating de novo prediction, homology-based prediction, and transcriptome-based prediction. De novo prediction involved using software such as AUGUSTUS (v.3.2.3)^59^, GENSCAN (v.1.0)^60^, GlimmerHMM (v.3.0.1)^61^, geneid (v.1.4)^62^, and NAP (v.2013.11.29)^63^ to predict coding regions from the genome with repetitive sequences masked. The homology-based prediction method downloaded protein sequence files of *Arabidopsis*, rice, *Panicum miliaceum*, *Panicum hallii*, and the published genome of a switchgrass line ‘AP13’ selected from the lowland ecotype ‘Alamo’ in the Phytozome database (https://phytozome-next.jgi.doe.gov/) and the National Center for Biotechnology Information (NCBI, https://www.ncbi.nlm.nih.gov/). These protein sequences were aligned to the two haplotype genomes of upland switchgrass using tblastN (v.2.2.26)^64^ with an e-value threshold of 1e^-5^. The Solar (v.0.9.6)^65^ software was used to integrate the BLAST results, and GeneWise (v.2.4.1)^66^ was employed to predict the precise gene structures in the corresponding genomic regions. The transcriptome-based prediction method used TopHat (v.2.0.13)^67^ and Cufflinks (v.2.1.1)^68^ to align transcriptome data to the two haplotype genomes. Trinity (v.2.1.1)^69^ was utilized to assemble RNA-seq data to create pseudo-expressed sequence tags (pseudo-ESTs), which were then mapped to the two haplotype genomes. Finally, EVidenceModeler (v.1.1.1)^70^ was used to integrate the gene sets obtained from the three methods into a non-redundant, more complete gene set (EVM sets). The Program to Assemble Spliced Alignments (PASA)^71^ was used to correct the EVM sets, adding information such as UTRs and alternative splicing, to obtain the final gene set.

### Annotation of protein-coding genes and non-coding RNA

Six databases were used for the functional annotation of coding genes, including Swiss-Prot (http://www.uniprot.org/)^72^, InterPro (https://www.ebi.ac.uk/interpro/)^73^, the Non-Redundant Protein Sequence database (NR, ftp://ftp.ncbi.nih.gov/blast/db/), the Pfam database (https://pfam-legacy.xfam.org/)^74^, the KEGG (http://www.genome.jp/kegg/)^75^, and the GO database (http://www.geneontology.org/page/go-database)^76^.

miRNA, rRNA, and snRNA were predicted in the genome using INFERNAL (v.1.1.5)^77^ with the Rfam database (https://rfam.org/)^78^. For tRNA, tRNAscan-SE (v.2.0.12)^79^ was used to predict tRNA sequences in the two haplotype genomes based on the structural characteristics of tRNA.

### Phylogenetic tree construction and divergence time estimation

BLASTP (v.2.7.1) was used to perform BLAST searches against the protein sequences from *P. hallii*, *Z. mays*, *S. bicolor*, and *O. sativa*, as well as the two subgenomes of ‘JJ31-B’ and ‘AP13’, with a default E-value of 1e^-5^. Orthofinder (v.2.3.1)^80^ with default parameters was then used to cluster the filtered BLAST results into paralogous and orthologous groups. The sequences of single-copy gene families were aligned using MUSCLE (v.3.8.31)^81^, and the alignment results were concatenated to form a super alignment matrix. RAxML (v.8.0.19; http://sco.h-its.org/exelixis/web/software/raxml/index.html)^82^ was used to construct the phylogenetic tree using the maximum likelihood method, with bootstrap values set to 100. The divergence time of each node on the phylogenetic tree was estimated using the MCMCTree program (v.4.5; http://abacus.gene.ucl.ac.uk/software/paml.html)^82^ with phylogenetic analysis by maximum likelihood (PAML) with the parameter settings ‘burn-in=10000, sample-number=100000, sample-frequency=2’. The TimeTree database (http://www.timetree.org/)^83^ provided species divergence times. On the basis of the orthologous genes for the two subgenomes each of ‘JJ31-B’ and ‘AP13’, the synonymous substitution (*Ks*) were calculated. The formula *t* = *Ks*/*2r* was used to estimate the divergence time between species, where r is the neutral substitution rate (r = 6.96 × 10^−9^)^28,84,85^.

### Identification of the *COR* gene families

The protein sequences of *Arabidopsis* and rice were downloaded from the TAIR (https://www.arabidopsis.org) and RGAP (http://rice.plantbiology.msu.edu) databases, respectively. Based on the 115 *COR* genes reported in *Arabidopsis*, we used BLASTP (v.2.7.1) to identify the COR protein sequences in rice, with an e-value set to 1e-10^29^. The top-ranked protein sequences were combined with the *Arabidopsis* protein sequences to create a merged library. Subsequently, we identified the COR protein sequences in ‘JJ31-B’ using an e-value of 1e-10 and identity > 60%^29^.

### Transcriptomic analyses of switchgrass under low temperature

Seeds of ‘JJ31’ and ‘Alamo’ were planted in plastic pots (10 × 15 × 6 cm) filled with quartz sand and placed in a growth chamber (26 °C with 14 hours of light, 22 °C with 10 hours of darkness). Cold stress treatment was then applied to E3 stage^40^ seedlings of both ecotypes, with conditions set to 4 °C with 14 hours of light and 4 °C with 10 hours of darkness, while the control group was maintained under normal conditions. After 12, 24, and 48 hours of cold stress treatment, the leaves and roots of JJ31 and Alamo were collected and stored at −80 °C. Three biological replicates were set for each treatment and control, with each replicate consisting of a mixture of three seedlings. RNA was extracted from the mixed samples using the RNeasy Plant Mini Kit (QIAGEN), and the quality of RNA was assessed by RNA gel electrophoresis. High-quality RNA was used to construct cDNA libraries with the NEBNext Ultra Directional RNA Library Prep Kit. Transcriptome sequencing was performed on the Illumina HiSeq X platform. The raw data were processed to remove adapters and low-quality nucleotide sequences using Trimmomatic (v.0.36)^41^. The quality of the filtered data was assessed using FastQC (v.0.11.9, https://www.bioinformatics.babraham.ac.uk/projects/fastqc/). A genome index file was built with Kallisto (v.0.46.0)^86^ using ‘JJ31-B’ as the reference genome.

Subsequently, the filtered transcriptome clean reads were aligned to the index file to obtain gene count values and transcripts per million (TPM). DESeq2 (v.1.24.0)^87^ was employed for identifying DEGs (|log_2_ (fold change)| ≥ 0.8 and adjusted *P*-value < 0.05) based on gene count values. GO and KEGG enrichment analyses were performed using the OmicShare tools (http://omicshare.com/tools).

### Physiological index measurement

Leaves of E3 stage^40^ seedlings of ‘JJ31’ and ‘Alamo’ were used for physiological measurements, with the cultivation methods and conditions being the same as those used for the seedlings prepared for transcriptome sequencing. The RWC, REC, and MDA content of the leaves were measured on seedlings after 1, 7, 14, 21, 28, and 35 days under both cold treatment and normal conditions. Transgenic rice and WT rice were cultivated for 45 d under 26°C with 14 hours of light and 22°C with 10 hours of darkness, followed by cold stress treatment at 4°C. After 24 h of cold stress, the MDA content and the activities of POD and SOD enzymes were quantified using rice leaves. The RWC of the leaves was determined using the saturated weighing method^88^ based on the formula *RWC* = (*FW*-*DW*)/(*TW*-*DW*), where *FW* refers to the fresh weight of leaves taken from the same part of seedlings, *TW* is the saturated fresh weight of these leaves after absorbing water, *DW* refers to the dry weight of leaves after soaking, blanching at 105 □ for 30 minutes, and then drying at 65 □ until a constant weight is reached. The measurements of REC, MDA, POD, and SOD were based on the methods previously described by our laboratory^42^.

### Differential expression analysis of allelic genes

Protein sequences from the two haplotype genomes were retrieved using TBtools (v.2.069)^89^. The proteins in ‘JJ31-A’ were compared to those in ‘JJ31-B’ using BLASTP (v.2.7.1), and syntenic blocks within the genomes were identified using MCScanX^90^ with default parameters. Finally, gene pairs with unique alignment relationships between the ‘JJ31-A’ and ‘JJ31-B’ genomes were obtained, with alleles required to originate from the same pair of homologous chromosomes. The sequences of ‘JJ31-A’ and ‘JJ31-B’ were combined into a single file^39^. An index file for the combined sequences was created using Kallisto (v.0.46.0)^86,91^, and the clean RNA-seq data were aligned to the index file to obtain gene count values. Pairwise comparisons of allelic genes (JJ31-A/JJ31-B) were performed using DESeq2 (v.1.24.0)^87^ based on the gene count values to identify differentially expressed genes, with criteria set at |log_2_(fold change)| ≥ 1 and adjusted *P*-value < 0.05. Genes meeting the following three conditions were identified as ASE genes: (1) the fold change of one allele compared to the other was > 2 or < 0.5; (2) TPM values > 1 in all transcriptome samples; (3) differential expression of alleles in at least one transcriptome sample. Transcriptome samples involved in ASE identification had at least two replicates.

### SNP calling

SNP calling was performed using GATK (v.4.3.0.0)^92^, with detection by HaplotypeCaller and genotyping *via* GenotypeGVCFs. The SelectVariants tool was used to obtain a collection of SNPs based on the "--select-type-to-include SNP" parameter. This collection was then filtered using the parameters "QD < 2.0 || FS > 60.0 || SOR > 3.0 || MQ < 40.0 || MQRankSum < −12.5." Finally, VCFtools (v.0.1.16)^93^ was employed to further filter the data using parameters with "--max-missing 0.9, --maf 0.05, --minDP 10."

### SV detection

To improve the accuracy of structural variant (SV) identification, we employed three tools: Manta (v.1.6.0)^94^, Delly (v.1.1.6)^95^, and LUMPY (v.0.3.1)^96^. First, we used LUMPY with the parameters -P -B -S -D to detect SVs, excluding insertions.We filtered results lacking split read support and conducted genotyping with SVTyper (v.0.7.1)^97^. The other two tools were used with their default settings for both SV detection and genotyping. Finally, we merged and filtered the results from these three tools using SURVIVOR (v.1.0.7)^98^, with the parameters set to “SURVIVOR merge 1000 3 1 1 0 50.” Only SVs identified by all three tools were retained.

### Selective sweep analysis

To identify genomic regions under selection in upland relative to lowland ecotypes, we used VCFtools (v.0.1.16)^93^ to perform *Fst* analysis based on a sliding window of 100 kb with a step size of 10 kb^99^. The top 5% windows were identified as selective sweeps^100^.

### Genome-wide association study

To improve the accuracy of GWAS results, we filtered the SNP and SV variant datasets, removing data with a minor allele frequency (MAF) < 0.05 or missing rate > 0.2. Association analysis was performed using GEMMA (v.0.94.1)^101^ based on a mixed linear model. The model is calculated as *y* = *Xα* + *Sβ* + *Kμ* + *e*, where *y* represents the phenotype, *X* represents the genotype, *S* is the population structure matrix, and *K* is the kinship matrix. *Xα* and *Sβ* represent fixed effects, while *Kμ* and *e* represent random effects.

### Transgenic rice validation

The CDS sequence of *PVB_3K03611.1* was synthesized using gene synthesis methods and inserted into the pCAMBIA3300-35S-EGFP vector under the control of the 35S promoter. The wild rice variety used for transgenic verification experiments in this study was Nipponbare (*O. sativa* L. spp. *japonica*). The transformation was performed using the *Agrobacterium*-mediated method as described by Hiei et al^102^. Firstly, *Agrobacterium* was added to the infection solution to prepare a resuspension with OD_600_ = 0.2. The rice callus was immersed in the *Agrobacterium* resuspension for 10-15 min, and then co-cultivated on medium at 20 °C for 48-72 h. The callus was then transferred to the selection medium and cultured in the dark at 26 °C for 20-30 d. The positive callus tissues screened were inoculated into the secondary screening medium and cultured at 26 □ in the dark for 7-10 d. The positive callus tissues that passed the secondary screening were inoculated into the differentiation medium and cultured at 25-27 □ in the light for 15-20 d. After the 2-5 cm buds appeared, they were inoculated into the rooting medium and cultured at 30 □ in the light for 7-10 d. PCR-positive seedlings were transplanted into soil and grown under conditions of 26 °C with 14 h of light and 22 °C with 10 h of darkness. When the plants reached the four-leaf stage, real-time quantitative PCR was performed, with each sample tested in three technical replicates. Primer information is provided in Supplementary Table 23.

